# Hook-length of the bacterial flagellum is optimized for maximal stability of the flagellar bundle

**DOI:** 10.1101/336602

**Authors:** Imke Spöring, Vincent A. Martinez, Christian Hotz, Jana Schwarz-Linek, Keara L. Grady, Josué M. Nava-Sedeño, Teun Vissers, Hanna M. Singer, Manfred Rohde, Carole Bourquin, Haralampos Hatzikirou, Wilson C. K. Poon, Yann S. Dufour, Marc Erhardt

## Abstract

Most bacteria swim in liquid environments by rotating one or several flagella. The long external filament of the flagellum is connected to a membrane-embedded basal-body by a flexible universal joint, the hook, which allows the transmission of motor torque to the filament. The length of the hook is controlled on a nanometer-scale by a sophisticated molecular ruler mechanism. However, why its length is stringently controlled has remained elusive. We engineered and studied a diverse set of hook-length variants of *Salmonella enterica*. Measurements of plate-assay motility, single-cell swimming speed and directional persistence in quasi 2D and population-averaged swimming speed and body angular velocity in 3D revealed that the motility performance is optimal around the wild type hook-length. We conclude that too short hooks may be too stiff to function as a junction and too long hooks may buckle and create instability in the flagellar bundle. Accordingly, peritrichously flagellated bacteria move most efficiently as the distance travelled per body rotation is maximal and body wobbling is minimized. Thus, our results suggest that the molecular ruler mechanism evolved to control flagellar hook growth to the optimal length consistent with efficient bundle formation. The hook-length control mechanism is therefore a prime example of how bacteria evolved elegant, but robust mechanisms to maximize their fitness under specific environmental constraints.

**Author summary:** Many bacteria use flagella for directed movement in liquid environments. The flexible hook connects the membrane-embedded basal-body of the flagellum to the long, external filament. Flagellar function relies on self-assembly processes that define or self-limit the lengths of major parts. The length of the hook is precisely controlled on a nanometer-scale by a molecular ruler mechanism. However, the physiological benefit of tight hook-length control remains unclear. Here, we show that the molecular ruler mechanism evolved to control the optimal length of the flagellar hook, which is consistent with efficient motility performance. These results highlight the evolutionary forces that enable flagellated bacteria to optimize their fitness in diverse environments and might have important implications for the design of swimming micro-robots.

## Introduction

Bacterial flagella are complex rotary nanomachines and enable directed movement of cells through various environments. The ability for locomotion and the presence of external flagellar structures contribute to pathogenesis and biofilm formation of many bacterial pathogens [1–4].

The bacterial flagellum is composed of three main structural parts: (i) a membrane-embedded basal body; (ii) a several micrometer long external filament; and (iii) the hook, a linking structure that connects the basal body and the rigid filament [5]. The basal body complex functions as a motor to rotate the flagellum and includes a proton motive force (pmf) dependent flagellum-specific protein export machine [6,7]. Assembly of the flagellum initiates with formation of the export machinery within the cytoplasmic membrane, followed by formation of a rod structure that traverses the periplasmic space. Assembly of the hook starts upon completion of the P- and L-rings, which form a pore in the outer membrane and polymerize around the distal rod [8,9]. The extracellular, flexible hook structure functions as a universal joint and allows the conversion of the torque generated by the cytoplasmic motor into rotational motion of the flagellar filament irrespective of the cell body orientation [10]. The self-assembly of the rod, hook and filament structures are controlled by different regulatory mechanisms. The length of the flagellar rod of ~25 nm is determined by the width of the periplasmic space [11], whereas the growth rate of flagellar filaments decreases with length and is controlled through pmf-dependent injection and diffusive movements of filament subunits inside the flagellar secretion channel [12]. The length of the hook structure is controlled to ~55 nm through a molecular ruler mechanism (Fig 1A) [13,14]. Upon termination of hook growth, the export apparatus switches secretion specificity from early (‘rod/hook’-type) to late (‘filament’-type) substrate secretion [15]. This mechanism ensures that a functional hook-basal-body complex is present, on top of which the long flagellar filament made of several tens of thousands flagellin molecules can assemble [16]. The FliK protein is responsible for both the measurement of hook-length and the transmission of the hook growth termination signal to the export apparatus, after which the switch in substrate specificity occurs. FliK functions as a molecular ruler, which takes intermittent length measurements throughout the assembly of the hook structure and terminates further hook growth when the hook has reached a length of ~55 nm or longer [14,17,18].

**Fig 1:**
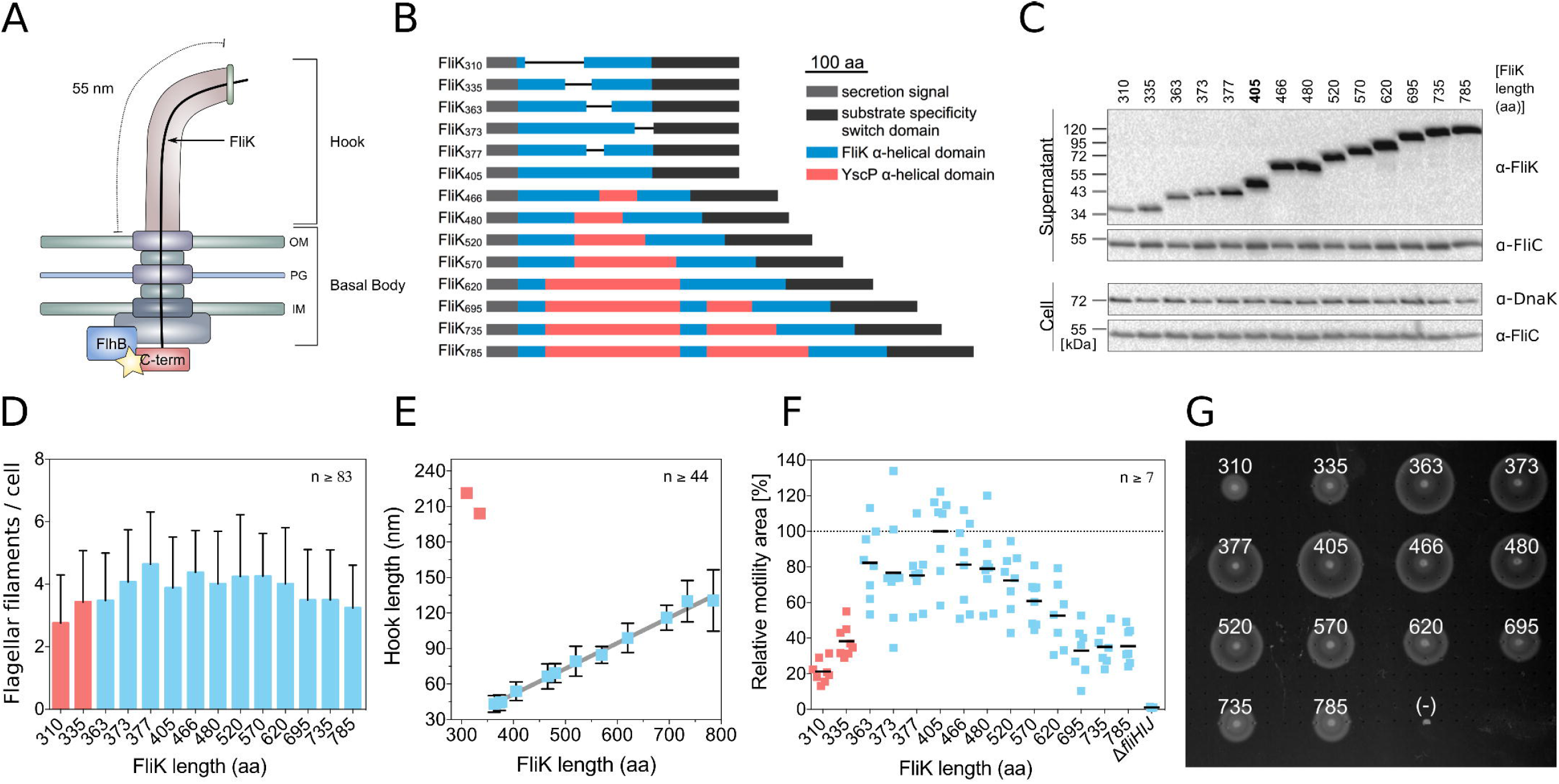
Characterization of hook-length variants and motility performance in semi-solid media. (A) Model of the flagellar hook-length control mechanism. The yellow star indicates the switch in substrate specificity within the export apparatus. (B) Engineered FliK length variants. (C) Cellular (cell) and secreted (supernatant) levels of FliK and flagellin (FliC). Loading control (DnaK). A representative gel of two independent experiments is shown. (D) Flagellation of hook-length variation mutants (mean number of flagella per cell + s.d.). FliK mutants with uncontrolled hook-length are highlighted in red. Representative fluorescent microscopy images are shown in Fig S2. A summary of statistical analysis of the number of flagellar filaments per cell is given in S1 Data. (E) Correlation between the length of the FliK molecular ruler and the hook-length of FliK mutants with controlled hook-length (mean hook-length ± s.d.; mean hook-length of the uncontrolled FliK mutants (red)). Representative electron micrographs are shown in Fig S3. (F) Quantification of swimming motility phenotype in semi-solid agar plates based on TB broth (1% tryptone and 0.5% NaCl) containing 0.3% agar (mean motility (black bars) normalized to the wt). Individual replicates are shown. (G) Representative motility swarms of a semi-solid agar plate of the data shown in panel F (FliK length in amino acids is indicated; non-flagellated control strain Δ*fliHIJ* (-)).

In peritrichously flagellated bacteria, the hook as a flexible linking structure allows formation of flagellar bundles at either cell pole irrespective of the relative position of the membrane-embedded motors. When all flagella turn counterclockwise (CCW), they form a bundle, causing the cell body to rotate in the opposite direction, and propel the cell forward (run phase). Additionally, the body has a secondary motion superimposed on the rotating axis called body wobbling due to the flagella bundle pushing the body off-axis [19, 20]. When one or more of the motors turn clockwise (CW), their associated flagella separate from the bundle, causing a random change in the cell orientation (tumble). After all the motors have reverted to CCW rotation, the flagellar bundle reforms and the bacterium swims in a new direction. In homogenous environment (absence of gradient), the run-and-tumble behavior results in an effective diffusive motion over large distances. Biasing the tumbling rate allows chemotaxis, *i.e*. seeking attractants or escaping repellents in a chemo-gradient [21–24].

The relationship between hook-length and motility is poorly studied. Loss of hook-length control diminishes motility [15]; however, the mechanisms have not yet been examined. The physiological benefit of tight hook-length control remains unclear and was highlighted recently as one of the big questions in bacterial hydrodynamics [25]. Our results reveal an optimal motility performance for a wild type hook-length of ~55 nm in respect to swimming speed, directional persistence and propulsive efficiency. We conclude that the molecular ruler mechanism evolved to control an optimal length of the hook structure for maximized motility performance via a more stable bundle formation. Thus, a mechanism to control flagellar hook-length optimizes the cell’s motility performance by minimizing random directional change during the run phase.

## Results

### FliK molecular ruler deletion and insertion variants are secreted and functional

The length of the flagellar hook is determined by secretion of a molecular ruler protein, FliK. We engineered FliK mutants varying from 310 to 785 amino acids (aa) length using insertions of α-helical parts of a homologous molecular ruler of the related virulence-associated type III secretion system of *Yersinia*, YscP, or deletions of the central, α-helical domain of FliK (Fig 1B) [14,17,26]. All FliK mutants were secreted and retained the ability to induce the switch in secretion specificity to late substrate secretion as evidenced by comparable levels of external flagellin FliC and induction of a Class 3 gene reporter (Fig 1C, S1 Fig). The flagellation levels of the FliK mutants were not significantly different with an average of 3.8 ± 1.5 flagella per cell (averaged across all FliK mutants) compared to 3.6 ± 1.3 flagella per cell for the wild type (wt) except for the shorter FliK variants FliK_310_ and FliK_377_ (Fig 1D, S2 Fig, S1 Data). We next purified hook basal body (HBB) complexes of the FliK length variation mutants and determined the length of the hook structures by transmission electron microscopy. FliK mutants ranging from 363 to 785 residues lengths retained a tight control of hook-length from 42 nm to 135 nm with a linear increase in hook-length of 0.2 nm per inserted amino acid (Fig 1E, S3 Fig). An increase of 0.2 nm per inserted amino acid is consistent with an alpha-helical conformation of the ruler domain of FliK, as suggested before [17]. The two shortest FliK mutants, FliK_310_ and FliK_335_, displayed partially uncontrolled hook lengths, indicating that a minimal FliK length is needed for effective hook-length control (Fig 1E, S3 Fig).

### Hook-length mutants display a pronounced motility defect in semi-solid medium

We next analyzed the set of hook-length variation mutants for their motility phenotype in semi-solid agar plates (Fig 1F, Fig 1G, S4 Fig). The motility halo size was substantially decreased for mutants with shorter or longer hooks (FliK ≤ 363 aa and FliK ≥ 520 aa) and peaked at hook lengths around the wt length of 55 nm (FliK = 405 aa). The observed motility defect for long and short hook mutants was independent of the motility buffer and agar concentration, while prolonged incubation highlighted the motility differences (S4 Fig, S5 Fig). These observations suggest that the hook-length of the wt is in some way optimal (Fig 1F, Fig 1G). We note, however, that the motility phenotype in semi-solid agar plates is a complex combination of bacterial growth, motility and chemotactic behavior [27,28]. We found that bacterial growth rate was not impaired in the FliK mutants (S5 Fig). Further, the presence of motility halos for all hook-length mutants indicated functional chemotaxis. However, in order to decouple chemotaxis from motility and to reveal the effect of hook-length on the swimming behavior, we next performed single-cell tracking experiments.

### Single-cell tracking reveals that hook-length mutants have lower swimming speed and shorter directional persistence

To identify which behavioral parameter explains the phenotype of hook-length mutants in semi-solid medium, we characterized single-cell behavior in a quasi 2D environment as described before [29]. In the absence of chemotactic signals, we observed the effect of hook-length variation on the basal run-and-tumble behavior. We found that cells with longer hook lengths (>75 nm; FliK > 520 aa) have reduced average swimming speed and tumble more frequently (Fig 2A, Fig 2B, S1 Movie). Consequently, the directional persistence and the effective diffusion coefficient [30] of the mutants are reduced (Fig 2C, Fig 2D). Therefore, longer hooks affect the cell’s ability to explore their environment. We propose that these observations are largely responsible for the motility phenotypes that we observed in semi-solid agar plates. We further note that the FliK variants FliK_363_ and FliK_373_, which produce hooks of slightly shorter length than the wt, displayed in liquid culture a motility behavior similar or slightly better than the wt. However, the shorter FliK variants performed poorer in semi-solid agar plates (Fig 1F) and we thus conclude that short hooks might have a decreased ability for productive tumble events due to increased stiffness and thus impair cell re-orientation in semi-solid agar and/or gradient environments.

**Fig 2:**
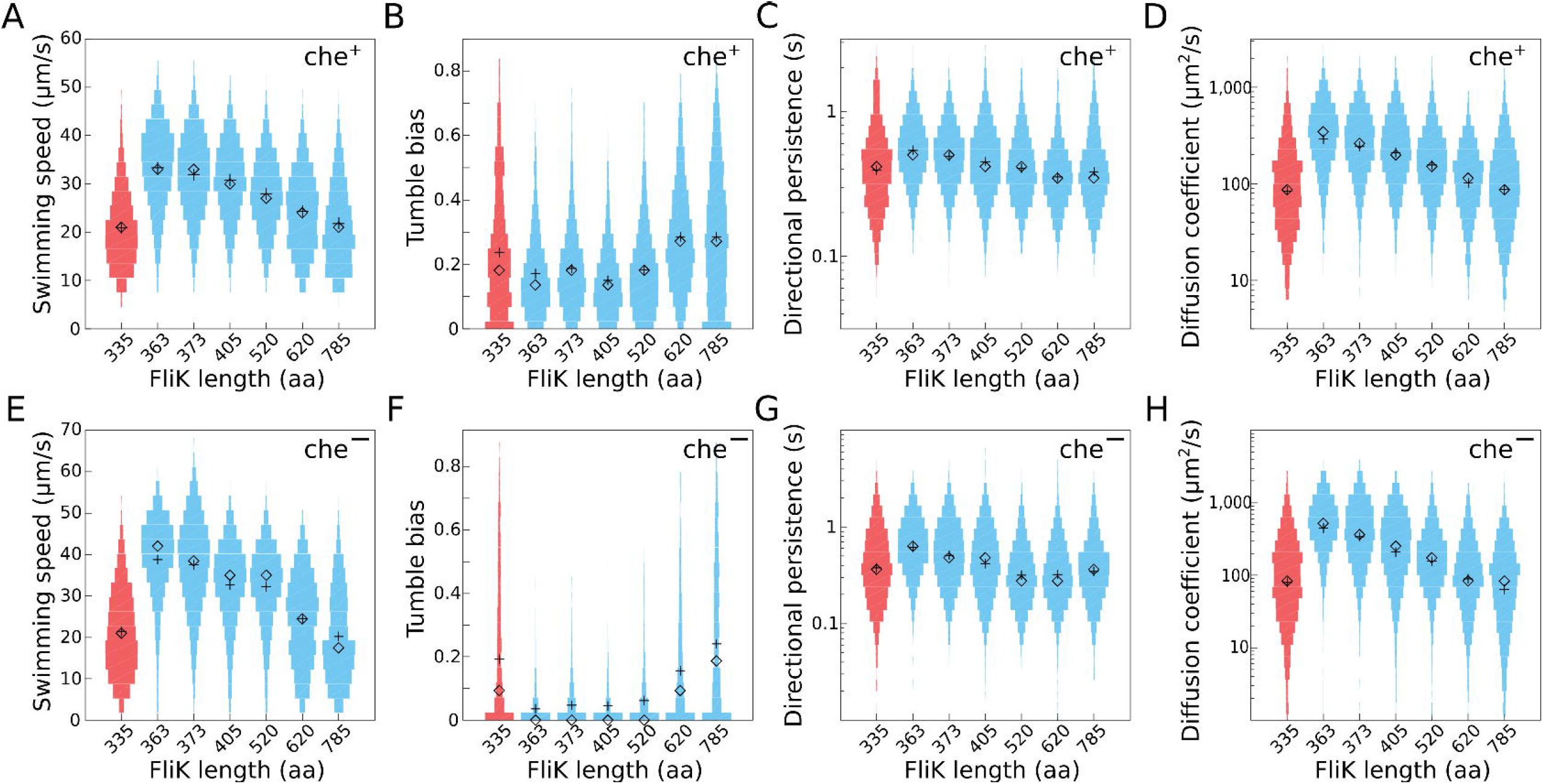
Single-cell motility performance of hook-length variants in quasi 2D. Distributions of single-cell motility parameters for different hook-length mutants in a wt background (che^+^). Distributions of (A) the average swimming speed of individual cells, (B) tumble bias (where tumbles are detected as a sudden change in direction regardless of the direction of the motor rotation), (C) directional persistence, (D) diffusion coefficient. (E-H) Same distributions from the smooth swimming hook-length mutants due to deletion of *cheY* (che^−^). Each distribution was calculated from the behavior of more than 600 individual cells pooled from three independent experiments. Mean: +, median: ◊. Uncontrolled hook-length mutants are shown in red.

We hypothesized that variations in the hook-length increase instability during formation of the flagellar bundle. Indeed, events in which filaments underwent polymorphic transformations without filaments leaving the bundle (by contrast with a single tight bundle), leading to small deflections in swimming direction, have previously been observed and speculatively associated with motor reversal [31]. To determine if the apparent increase in tumble bias is the result of an increased rate of polymorphic changes in the bundle, which are not associated with reversions of the flagellar motor rotation, we characterized the behavior of same hook mutants in non-chemotactic, smooth-swimming cells (Δ*cheY*). Mutants lacking CheY do not tumble and thus do not respond to chemical gradients and do not efficiently navigate the agar gel matrix of semi-solid agar plates [27]. Accordingly, we assessed the motility behavior of smooth-swimming hook-length mutants using 2D single-cell tracking. In the Δ*cheY* genetic background (che^−^), hook-length mutants still have reduced swimming speeds and smaller diffusion coefficients when compared to the wt hook-length (Fig 2E, Fig 2H). Cells with hooks longer than 75 nm have a higher probability of changing direction suddenly; events, which we will refer to as ‘pseudo-tumbles’ (Fig 2F). In non-chemotactic, smooth-swimming Δ*cheY* mutants, these ‘pseudo-tumbles’ cannot be caused by reversions of the flagellar motor.

Finally, che^−^ hook-length mutants displayed a reduced directional persistence (Fig 2G), because of a general increase in the probability of large changes in swimming direction (S6 Fig). These observations suggest that the poorer swimming performance in hook-length mutants is likely to be primarily due to decreased stability of the flagellar bundle. To further analyze the dependence of ‘pseudo-tumble’ events on flagellar hook-length and to estimate a tumbling rate, we calculated the directional autocorrelation function and mean square displacement (MSD) of selected non-chemotactic (ΔcheY), smooth-swimming hook-length mutant from the single-cell tracking experiments (S7 Fig, S1 Text). The autocorrelation function measures the persistence of bacterial motion and the MSD represents the spatial exploratory potential of each cell. The observed trajectories of non-chemotactic, hook-length mutants with a ‘pseudo-tumble’ phenotype were fitted to a simple run-and-tumble migration model (S2 Text, S8 Fig), taking into account the curvature of the tracks due to hydrodynamic interactions of bacteria swimming in quasi 2D close to the surface. The fit of the run-and-tumble model to the trajectories of the non-chemotactic, hook-length mutants suggest that the directional persistence decreases with increasing hook-length (Fig 2G, S7 Fig). We concluded that the pseudo tumble events of the non-chemotactic mutants arise from a mechanical instability of the hook, similar to the tumbling mechanism of *V. alginolyticus* [32].

### Non-chemotactic long hook mutants display a pseudo-tumble behavior

The conclusion we have reached leads to an interesting prediction of the behavior of hook mutants in agar gels. Chemotactic (che^+^) bacteria can navigate through an agar gel matrix due to their run-and-tumble motility behavior (Fig 3A) [27]. However, non-chemotactic, smooth-swimming (Δ*cheY*) cells are trapped because tumbling is required to efficiently escape the agar gel matrix (Fig 3B). Accordingly, if the increased flagellar bundle instability in hook mutants conferred on them a ‘pseudo-tumble’ phenotype, then we may expect che^−^ long hook mutants to regain the ability to migrate through an agar gel matrix.

**Fig 3:**
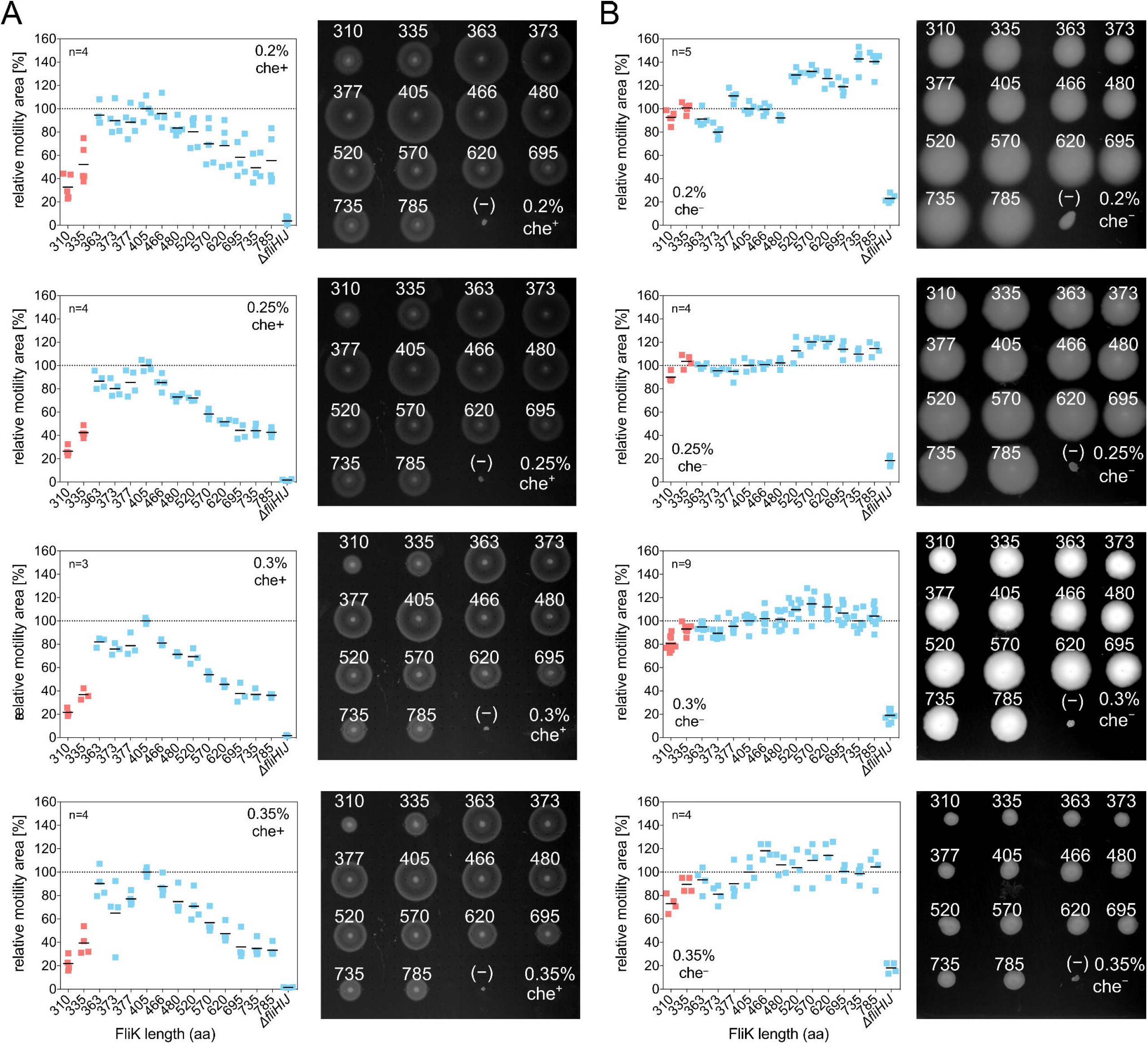
Motility behavior of chemotactic and non-chemotactic hook-length variants in semi-solid agar. (A) Motility performance of chemotactic (che+) and (B) non-chemotactic (Δ*cheY*, che^−^) hook-length mutants in semi-solid agar plates based on TB broth (1% tryptone and 0.5% NaCl) with varying agar concentrations (0.2%, 0.25%, 0.3%, 0.35% agar). The che+ mutants were incubated for 3-4 h whereas the che^−^ mutants were incubated for 14-18 h, depending on the agar concentration. For all panels, a representative motility plate is shown on the right-hand side. FliK length in amino acids is indicated; (-) annotates the non-flagellated control strain Δ*fliHIJ*. Individual replicates are shown and black bars represent the mean motility normalized to the wt. FliK mutants with uncontrolled hook-length are highlighted in red.

A similar pseudotaxis behavior has previously been observed for non-chemotactic *E. coli* mutants, where point mutations in the switch complex proteins FliG and FliM allowed random motor rotational switching in the absence of chemotactic stimuli [27], as well as in a non-chemotactic *A. tumefaciens* strain, where suppressor mutations in the hook, the *fliK* gene and the motor force generators were isolated that allowed the che^−^ cells to navigate through semi-solid agar [33].

We analyzed the motility behavior of non-chemotactic, smooth-swimming hook-length mutants to test our prediction. The motility halo size of che+ hook-length mutants peaked around the wt hook-length (Fig 1F, Fig 3A). In contrast, che^−^ long hook mutants performed substantially better than che^−^ mutants with shorter and wt hook-length (Fig 3B), in accordance with our suggestion that long hook lengths confer a pseudo-tumble phenotype. We also noticed a big difference in the speed of spreading, che^−^ mutants needed to be incubated ~5× longer to reach comparable halo sizes. This indicates that pseudo-tumbling events occur less frequently than tumbling, *i.e*. due to motor reversal. There is an interesting difference in appearance, che^−^ colonies form disks rather than typical chemotactic rings confirming the pseudo-tumble events are not related to chemotaxis and thus not associated with motor reversal.

### Motility parameters are optimal for wt hook-length in 3D liquid environment

In peritrichously flagellated bacteria, CCW rotation of all flagella allows formation a flagella bundle rotating at speed ω and results in a run phase with the cell moving forward at speed *v*. The flagella bundle further pushes the body off-axis and results in body wobbling, which is characterized by an angular velocity **Ω** [19, 20]. For flagellated bacteria swimming in a Newtonian liquid, *e.g*. buffer, *v* is proportional to **Ω** and ω, which are determined by the geometry of both body and flagellar bundle [34].

All the experiments described so far pertain to bacteria swimming close to a hard surface or in a gel matrix. In a final set of experiments, we measured swimming speed *v* and body angular velocity Ω using high-throughput DDM and DFM in a 3D liquid environment [34–36]. DDM and DFM experiments were performed with bacteria grown in TB medium at 30 °C, which did not affect the overall motility behavior (S9 Fig, S3 Text). We recorded over 10^4^ cells swimming in a 3D liquid medium (i.e., in bulk) for each of the hook-length mutants. We found that the population-averaged swimming speed *v* was maximal whereas the population-averaged body angular velocity Ω was minimal over the range FliK ≥ 363 aa and FliK ≤ 480 aa, encompassing the wt (FliK = 405 aa) (Fig 4A, Fig 4B, S10A Fig, S10B Fig). Some variation in *v* and **Ω** were observed between independent datasets, but this disappeared when we calculated the population-averaged propulsive efficiency (processivity) 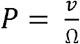, which is the average run length per revolution. We found a clear maximum in processivity near the wt hook-length, in the range 363 aa ≤ FliK ≤ 405 aa (Fig 4C, S10C Fig). Moreover, the swimming speed distribution was narrower (the width of the speed distribution *S* is minimal) in the range 363 aa FliK 570 aa (S11 Fig), and swimming trajectory appeared to be straighter (*R*, the ratio of two speeds measured at two length scales, is minimal; see Methods for more details) in the range 405 aa ≤ FliK ≤ 520 aa. Variations in 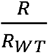 (with 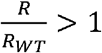) with FliK length are indicative of decreased directional persistence in the cell trajectories (Fig 4D, S10D Fig, S10E Fig). The increase in cell body angular velocity and the resulting decrease in processivity support the notion that deviations from the wt hook-length increase instability in the flagellar bundle and affect both swimming speed and directional persistence.

**Fig 4:**
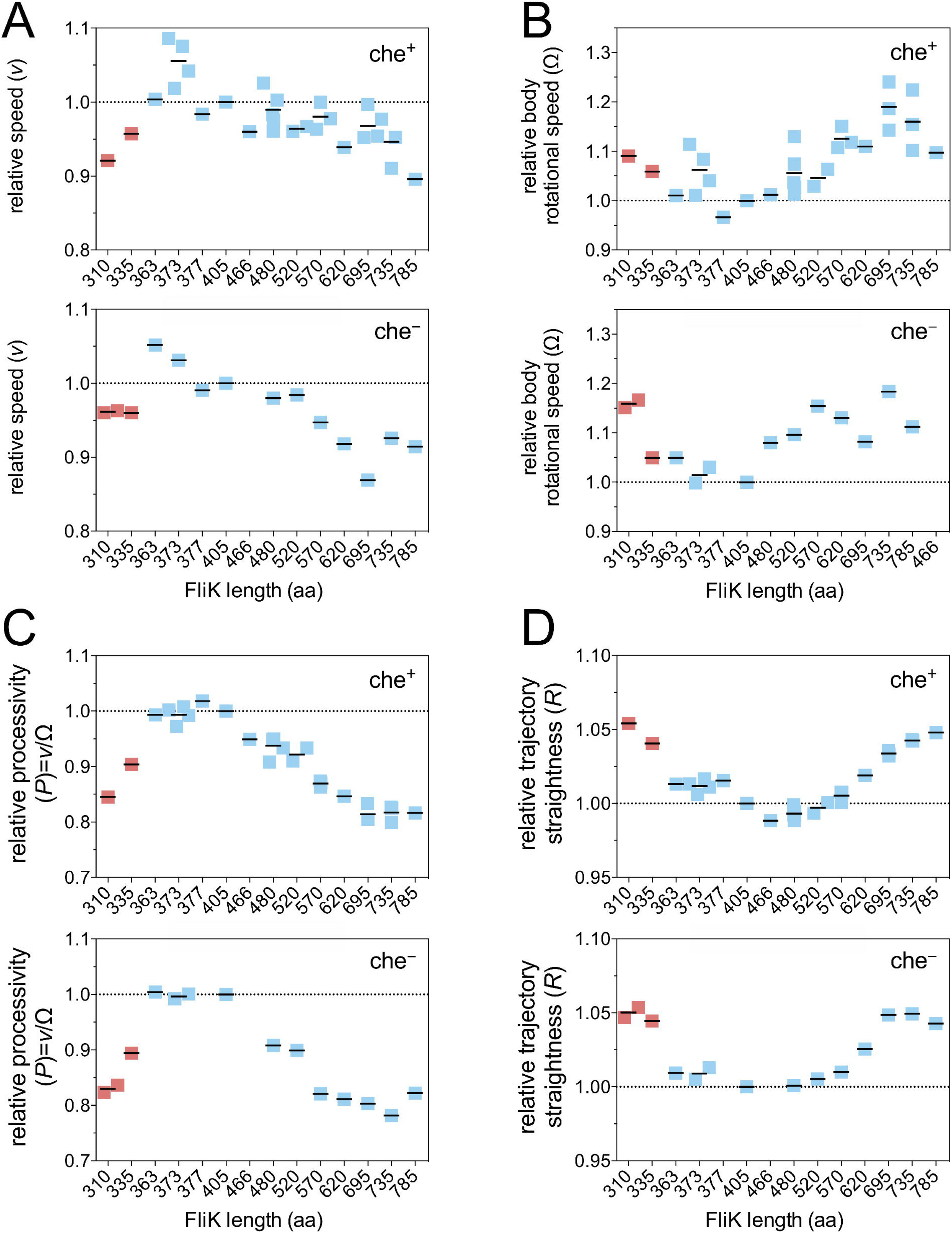
Motility performance of hook-length mutants in 3D. (A) Swimming speed *v*, (B) body rotational speed **Ω**, (C) processivity P=*v*/**Ω**, (D) straightness of trajectory *R*. Data points represent individual experiments normalized to the wt (black bars; mean). Uncontrolled hook-length mutants are shown in red. All parameters were obtained by normalizing the time-dependent parameters to the wt (FliK405) and by averaging over the time-window 10 min<t<60 min.

These experimental findings are independent of the tumbling behavior, as run-and-tumble strains displayed similar hook-length dependency compared to the smooth-swimmers (Fig 4, S12 Fig). This was expected because the motility parameters are measured over a length-scale corresponding to the run length (<15 μm, distance between two tumble events). Differences are found in absolute values of *R* between run-and-tumble and smooth-swimmers at a given hook-length due to tumbling events (S12D Fig). However, the relative path straightness 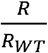 showed the same behavior for che^+^ and che^−^ hook mutants (Fig 4, S12 Fig).

## Discussion

In summary, our results provide substantial evidence that hook-length control in peritrichously flagellated bacteria has been evolutionary selected to optimize the stability of the flagellar bundle and thus maximize swimming performance. We found that swimming speed and directional persistence of the trajectories of swimming bacteria are tightly related to the length of the hook structure. Recently, small polymorphic changes in the flagellar bundle have been observed without filaments leaving the bundle. These polymorphic changes create deflection in the swimming trajectory and have been associated speculatively with motor reversal [31]. Our present work suggests these events might be associated with the run phase rather than motor reversal as they also occur in smooth swimming mutants. Thus, the role of hook-length control may be to minimize such polymorphic changes by maximizing stability of the flagellar bundle during the run phase. Accordingly, we propose that the hook-length control mechanism evolved to match the requirements for effective formation of the flagellar bundle. Stiffness is an inverse function of the hook-length, *e.g*. stiffness increases with decreasing hook-length. Too short hooks are too stiff to function as a universal joint whereas too long hooks may buckle and create too much instability, thus necessitating the need to control for an optimal length of the hook structure. Further, the observed higher processivity of hooks with wt length appears to result in more efficient propulsion *i.e*. the cells are able to achieve higher swimming speeds for the same energy expenditure. We further note that the mechanical instability of the hook might indicate polymorphic transitions of the hook [37] and, accordingly, the hook-filament flexibility could explain both the observed polymorphic changes in the flagellar bundle and instability.

Our results are also consistent with previous observations, where the wt hook of *E. coli* was modified to be twice more rigid by binding of streptavidin and the flagellar bundle was unable to form [38]. Hook-length control may also play a major role in other bacteria. Son and co-workers demonstrated for the polar flagellated *Vibrio alginolyticus* that a buckling instability of the hook enables the cell to ‘flick’ and re-orient [32]. They suggested buckling occurs when the viscous loads, *i.e*. force and torque exerted on the hook by the cell body and flagellum, exceed the hook’s buckling critical threshold. The associated critical torque and force are inversely proportional to the hook-length and its square, respectively. Thus, we expect that changing the hook-length of polar flagellated bacteria could dramatically alter the instability region and enable more “flicking” events, which in turn would affect motility and chemotactic behavior.

The function of the bacterial flagellum relies on the self-assembly of major components with defined or self-limiting lengths [11,12]. Incomplete assembly or small deviations from the blueprint result in a non-functional motility organelle with devastating consequences for the organism. Thus, regulatory mechanisms evolved to determine the correct dimensions of flagellar sub-assemblies and to ensure robust assembly of a functional motility organelle. Our results demonstrate the selective advantage of a mechanism that controls the length of the hook structure. In particular, our findings suggest that the molecular ruler mechanism evolved to determine the optimal length of the flagellar hook. These results highlight the evolutionary forces that enable flagellated bacteria to optimize their motility performance and fitness in diverse environments.

## Materials and Methods

### Bacterial strains, plasmids and media

All bacterial strains used in this study are listed in S1 Table. Cells were grown in either minimal medium containing 2% yeast extract (Vogel-Bonner medium E, 2% yeast extract), lysogeny broth (LB, 1% tryptone, 0.5% yeast extract, 0.5% NaCl) [39] or TB broth (1% tryptone and 0.5% NaCl) at 30 °C or 37 °C. The generalized transducing phage of *Salmonella enterica* serovar Typhimurium P22 *HT105/1 int-201* was used in all transductional crosses [40]. Insertions and deletions in the *fliK* gene were constructed using λ-RED homologous recombination [41].

### Growth curve

FliC-locked bacteria were grown overnight (o/n) at 37 °C and diluted to OD_600_ of 0.001 in LB (final volume 200 μl) in a honeycomb multiwall plate. The bacteria were incubated in the Bioscreen reader (EZExperiment software) at 37 °C with shaking and OD_600_ was measured every 15 min and the measurements were blank corrected.

### Motility Assay

Motility of the bacteria was assessed using semi-solid agar plates containing a range of agar concentrations from 0.2% – 0.35% essentially as described before [42]. Semi-solid agar plates were based on either minimal medium containing 2% yeast extract (Vogel-Bonner medium E, 2% yeast extract) or TB broth (1% tryptone and 0.5% NaCl). Equal amounts of overnight cultures were inoculated into the semi-solid agar plates using a pin tool and incubated at 37 °C for 3 – 4 h (che+) and 14 – 18 h (che^−^), respectively. The areas of the swimming halos (che^+^) or disks (che^−^) were analyzed using ImageJ and normalized to the wt (hook-length ~55 nm; FliK = 405 aa).

### SDS-PAGE and Western blot

FliC-phase locked bacteria were grown until late exponential phase in LB. 2 ml of culture was used for harvesting cells (whole cell lysate) and the supernatant by centrifugation (13,000 g, 4 °C, 5 min). Proteins were precipitated using 10% TCA and the samples were resuspended in SDS sample buffer normalized to their OD_600_. Proteins were separated by SDS PAGE and Western blot was performed using anti-FliK, anti-FliC and anti-DnaK antibodies.

### Hook-basal-body purification

Purification of hook-basal body complexes without C-ring was performed as described [43] with slight modifications. Briefly, bacteria were grown in 500 ml culture until OD_600_ 1 – 1.5. Cells were harvested (8,000g, 4 °C, 10 min) and resuspended in 30 ml ice-cold sucrose solution (0.5 M sucrose, 0.1 M Tris-HCl, pH 8). 3 ml lysozyme (10 mg/ml in sucrose solution) and 3 ml 0.1 M EDTA (pH 8) were added to digest the peptidoglycan layer and to prepare the spheroplasts. The suspension was stirred for 30 min on ice. 3 ml 10% Triton-X and 3 ml 0.1M MgSO_4_ were added to lyse the spheroplasts. For complete lysis of the spheroplasts the mixture was stirred overnight at 4 °C. Afterwards 3 ml 0.1M EDTA (pH 11) was added. Unlysed cells and cell debris was pelleted by centrifugation (17,000 g, 4 °C, 20 min) and the pH was adjusted to pH 11 by adding 5 N NaOH. Following centrifugation (17,000g, 4 °C, 20 min) the flagella were pelleted by ultracentrifugation (100,000 g, 4 °C, 1 h). The pellet was resuspended in 35 ml ice-cold pH 11 buffer (10% sucrose, 0.1 M KCl, 0.1% Triton-X) and centrifuged (17,000 g, 4 °C, 20 min) and the flagella were pelleted by ultracentrifugation (100,000 g, 4 °C, 1 h). The pellet was resuspended in 35 ml ice-cold TET-solution (10 mM Tris-HCl pH 8, 5 mM EDTA, 0.1% Triton-X) and the mixture again subjected to ultracentrifugation (100,000 g, 4 °C, 1 h). For depolymerization of the filaments, the flagella were resuspended in 35 ml pH 2.5 buffer (50 mM glycine, 0.1% Triton-X) and left at room temperature (RT) for 30 min. The debris was pelleted by centrifugation (17,000 g, 4 °C, 20 min) and the HBB were collected by ultracentrifugation (100,000 g, 4 °C, 1 h). The pellet was air-dried and resuspended in 200 μl TE solution (10 mM Tris-HCl pH 8, 5 mM EDTA). The samples were stored at 4 °C and imaged the following day by negative staining electron microscopy.

### Fluorescent microscopy

FliC-phase locked bacteria were grown until mid-log phase in LB or TB medium and immobilized on pre-coated 0.1% poly-L-lysine coverslips in a flow cell as described [14]. Cells were fixed with formaldehyde (2% final) and glutaraldehyde (0.2% final). Flagella were stained using polyclonal FliC antibody (anti-rabbit) and anti-rabbit coupled to Alexa Fluor 488. The membrane was stained using *N*-3-Triethylammoniumpropyl-4-6-4-Diethylamino Phenyl Hexatrienyl Pyridinium Dibromide (FM 4-64) and the DNA using 4’,6-diamidino-2-phenylindole (DAPI). Images were taken using a Zeiss AxioObserver microscope at 100x magnification.

### Electron microscopy

Hook samples were stored at 4 °C to straighten the hooks and facilitate length measurements. The samples were negatively stained by 2% aqueous uranyl acetate on a carbon film. Samples were imaged using a Zeiss TEM 910 at an acceleration voltage of 80 kV with calibrated magnifications. Images were recorded using a Slow-Scan CCD camera (ProScan, 1024×1024, Scheuring, Germany) with ITEM-Software (Olympus Soft Imaging Solutions, Münster, Germany). The length of the hooks was measured using ImageJ.

### Luciferase assay

The luciferase assay was performed as described before [18]. The plasmid pRG19 (*motA::luxCDABE*) was used for monitoring the switch from early substrate secretion to late substrate secretion. The class 3 gene *motA* is expressed only upon HBB completion. Synchronization of flagellar gene expression was achieved by placing the AnTc inducible *tetA* promotor upstream of the *flhDC* gene [44]. Cells were grown in white clear-bottom 96-well plates (Greiner) and production of light and the OD_600_ was measured over time using a Varioskan Flash multiplate reader (Thermo Fisher). After blank correction, relative light units (RLU) were calculated as:

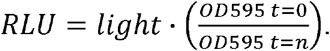

### 2D single-cell tracking and behavioral analysis

Overnight cultures were incubated in a shaking (200 rpm) incubator at 37 °C in LB broth. Overnight cultures were diluted 1:100 into fresh LB medium and incubated shaking at 200 rpm at 37 °C for 2.5 h. After cells reached the mid-exponential growth phase (OD_600_ ~0.5) cultures were diluted 1:50 in fresh LB medium and 7 μl of the dilution were placed on a microscope slide, covered with a #1.5 22×22 mm coverslip, and sealed with VALAP (equal amount of petrolatum, lanolin, and paraffin wax). The trajectories of swimming cells were recorded as previously described [29]. Briefly, cell positions were recorded using time-lapse phase contrast microscopy (Nikon Eclipse TI-E with 10× Nikon CFI Plan Fluor, N.A. 0.30, W.D 16.0 mm) at 10 frames per seconds (fps) (Zyla 4.2, Andor Technology Ltd.) for 100 seconds. The field of view contained ~200 cells on average over 1.3 mm^2^. Experiments were done in triplicate with independent biological samples. Individual trajectories were reconstructed as previously described using custom MATLAB (The MathWorks, Inc.) script (https://github.com/dufourya/SwimTracker) [29] and u-track 2.1 [45]. Tumbles were detected using a two-state Markov-chain model representing the ‘swimming’ and ‘tumbling’ states with bi-variate Gaussian distributions of the relative acceleration and angular acceleration. The Markov-chain model was trained on a reference dataset of wt *S. enterica* cell trajectories. The diffusion coefficient of individual trajectories was calculated using a covariance-based estimator [30] averaged over the time scales exceeding the time scale of rotational diffusion (>10 s), over which cell trajectories are diffusive. The directional persistence τ for each trajectory was calculated according to the relationship 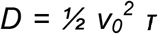, where *D* is the diffusion coefficient and *v_0_* is the average cell speed.

### 3D motility analyses using DDM and DFM

For DDM/DFM experiments of bacterial swimming behavior in 3D, cells are grown and washed, then loaded in sealed capillaries, placed onto a microscope, and movies were recorded. Importantly, these five steps are performed in parallel for a range of hook mutants, which allows precise measurement of potential small difference in motility. Details are given below.

As standard protocols, overnight cultures were grown in LB for ~16 h using a shaking incubator (200 rpm) at 30 °C. The overnight culture was diluted 1:100 in 35 ml TB and grown for 2.5 – 3 h at 30 °C with shaking (200 rpm) to OD_600_ ~0.4 – 0.5. In some cases (S9 Fig), overnight cultures were grown in LB for ~16 h using a shaking incubator (200 rpm) at 37 °C. The overnight culture was diluted 1:100 in 35 ml LB and grown for 1.5 – 2 h at 37 °C with shaking (200 rpm) to OD_600_ ~0.4 – 0.5.

Cells were washed three times with motility buffer (MB, pH = 7.0, 6.2 mM K_2_HPO_4_, 3.8 mM KH_2_PO_4_, 67 mM NaCl, and 0.1 mM EDTA) by careful centrifugation to minimize flagellar damage and were resuspended in MB to OD_600_ of 0.3. The washing step is performed simultaneously for several samples and takes typically ~30 min for 6 samples. We note that the viscosity of MB is similar to that of pure water with ~10^−3^ Pa · s [34].

Commercial glass capillaries (Vitrocom) with dimensions (L=5 cm x W=8 mm x H=400 μm, volume ~150 μl) were filled, sealed with petroleum jelly, and placed onto a fully-automated inverted microscope (Nikon TE300 Eclipse), and time-series of both phase-contrast (40 s long, Nikon Plan fluor 10× Ph1-NA=0.3 objective, Ph1 phase ring, 4,000 images at 512 × 152, 100 fps) and dark-field (10 s long, Nikon Plan Fluor 10× Ph1-NA=0.3 objective, Ph3 phase ring, 2,000 images at 512 × 152, 200 fps) movies are consecutively recorded continuously for ~1 h using an Hamamatsu ORCA-Flash 4.0 camera. One capillary at a time was prepared within 1 – 2 min, the movie was recorded immediately and then proceeded with next sample. Once all sample were prepared, continuous recording started. For 6 samples, this results in movie recorded every ~5 min for each capillary. Movies were recorded in bulk conditions (i.e. 100 μm above inner glass wall) to avoid interactions with wall.

From DDM analysis of the phase-contrast movies, the mean *v* and width *σ* (and consequently the relative width S = *σ* / v) of the swimming speed distribution *P*(*v*), averaged over 10^4^ cells, were extracted. Details can be found elsewhere [35, 36, 46] and only an overview is given below. Based on low-optical resolution, DDM characterizes the spatio-temporal fluctuations of the cell density via the so-called temporal autocorrelation function of the *q*-th Fourier component of the number density fluctuations, *f*(*q,τ*), with *τ* the delay time. *q* is the spatial frequency and defines a characteristic length-scale *L* = 2π / *q*. Given a suitable model for *f*(*q,τ*), kinetic parameters can be extracted. In case of swimming bacteria, DDM allows the measurement of an advective speed over a range of *q* values or length-scale *L*. The mean swimming speed *v* and width σ are extracted by averaging over the range 0.5 ≤ *q* ≤ 2, corresponding to 3 ≤ *L* ≤ 13 μm, and over the time window 10 min < t < 60 min.

A qualitative measure of the path straightness can also be estimated from DDM by calculating the ratio of speeds measured at two different length-scales, *R* = *v*(*L1*) / *v*(*L2*), with e.g. *L1* < *L2*. Theoretically, swimmers with a straight path yield *R* = 1. Deviations from straight-path, due to *e.g*. rotational diffusion or tumbling events, will increase this ratio above 1.

From DFM analysis of the dark-field movies, the body angular velocity **Ω** averaged over ~10^4^ cells in a few seconds was extracted. Details can be found elsewhere [34]. Briefly, the power spectrum of the flickering image of individual cells was Fourier transformed and the lowest frequency peak in the average power spectrum was identified as **Ω** / 2 π.

The numerical data used in all figures are included in S2 Data, S3 Data and S4 Data.

## Acknowledgments

We thank Nadine Körner for expert technical, Kelly ⊤. Hughes for generous donation of strains and Howard Berg for discussions.

## Supplementary Materials Legends

**S1 Text**: Observations from modeling of single-cell behavior.

**S2 Text**: Derivation of the velocity autocorrelation function for the circular run-and-tumble model.

**S3 Text**: Effects of growth medium and temperature on single-cell behavior.

**S1 Table**: List of *Salmonella* Typhimurium strains used in this study.

**S1 Fig: Substrate secretion switch of the hook-length mutants**. Expression of flagella synthesis was synchronized by induction of the P*_tetA_-flhDC* flagellar master regulatory operon. The switch in secretion specificity was monitored over time by measuring promoter activity of a Class 3 P*_motA_-luxCDABE* reporter. Individual panels compare a hook-length mutant (FliK length indicated in aa), the wt (FliK405) and the negative control (Δ*fliHIJ*). Uncontrolled hook-length mutants are shown in red.

**S2 Fig: Hook mutants express the same number of flagella**. (A) Quantification of the number of flagella of individual bacteria grown in LB at 37 °C. The flagellation pattern was determined by anti-flagellin immunostaining. Per mutant, the flagella per cell of all bacteria in 10 separate fields of view were counted manually. The average number of flagella per cell ± s.d. was calculated using Gaussian non-linear regression analysis. (B) Representative fluorescent microscopy images of anti-flagellin immunostaining. The flagellar filament was detected using anti-FliC immunostaining (green), membranes were stained using FM-64 (red) and DNA was stained using DAPI (blue). Scale bar = 5 μm.

**S3 Fig: Hook-length corresponds with FliK protein length**. (A) Histograms of hooks isolated from the FliK mutants. Hook-basal-body complexes were purified and imaged by negative staining electron microscopy. The average hook-length ± s.d. was calculated using Gaussian non-linear regression analysis. (B) Representative electron micrograph images of purified hook-basal-body complexes. Scale bar = 100 nm.

**S4 Fig: Swimming motility plate behavior of hook-length variants**. (A) Motility assay of the hook-length mutants in semi-solid agar plates based on minimal medium containing 2% yeast extract (Vogel-Bonner medium E, 2% yeast extract (YE)) and 0.3% agar. Quantification of the swimming motility assay (left panel) and representative semi-solid agar plate (right panel). FliK length in amino acids is indicated; (-) indicates the non-flagellated control strain Δ*fliHJ*. Motility halo size was measured using ImageJ and normalized to the wt (dots represent single data points; bars represent the mean; red dots represent the uncontrolled hook-length mutants). (B) Time course motility assay of the hook-length mutants in semi-solid agar plates based on TB broth (1% tryptone and 0.5% NaCl) and containing 0.3% agar. Motility halo size was determined every 30 minutes and the average motility halo area of 5 biological replicates is shown. Uncontrolled hook-length mutants are shown in red.

**S5 Fig: Growth analysis of the hook-length variants**. Bacteria were cultured in liquid LB medium and the OD_600_ was measured every 15 minutes. The average OD_600_ of 6 biological replicates is shown. Uncontrolled hook-length mutants are shown in red.

**S6 Fig: Motility performance of hook-length variants in quasi 2D**. The distributions of motility parameters for different hook-length mutants. Distributions of (A) the instantaneous speeds and (B) the angular velocities pooled from all the trajectories from cells with a wt genetic background (che+). (C-D) The same distributions from smooth swimming mutants due to deletion of *cheY* (che−). Each distribution was calculated from more than 1,000 minutes of cumulative trajectory times pooled from three independent biological replicates each containing more than 600 individual cells. Mean: +, median: ◊.

**S7 Fig: Autocorrelation functions and mean square displacement fits of the circular run-and-tumble model for selected hook-length mutants**. Observed orientational autocorrelation functions and circular run-and-tumble model autocorrelation function fits (A) and observed mean square displacements and circular run-and-tumble model mean square displacement fits (B) of chemotaxis deficient mutants (Δ*cheY*). Wt (FliK405) hook-length is shown at the top. Hook-length increases from top to bottom. The parameters of the model in each case are those reported in the supplementary text (S2 Text).

**S8 Fig: Circular run-and-tumble model**. (A) In this model, individual cells are assumed to move on a circle of radius *s* with an angular velocity *Ψ*. Under these conditions, the autocorrelation will be given by 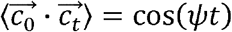. (B) and the mean square displacement will be given by 〈*r*^2^〉 = 2*s*^2^[1 − (*ψt*)] (C). (D) Autocorrelation function and (E) mean square displacement under the assumption that the angular velocities are distributed among the population according to *P*(*Ψ*). Shown are the observables when *P*(*ψ*) = *λe*^−*λ*^. (F) Sample track of a virtual cell in the circular run-and-tumble model. The cell starts moving on a circle as in the uniform circular motion model; however, at exponentially distributed waiting times, the cell will reorient (tumble) and move along a new circular path. An example of autocorrelation of a single realization of the model is shown in panel G. An example of square displacement of a single realization of the model is shown in panel H. The value of the circular path radius *s*, necessary for the calculation of (*r*^2^) was obtained with the relation between the angular velocity ψ and tangential velocity *v*, ψ = *v*/*s*. The value of *Ψ* was obtained from the fitting of the model to the experimentally observed autocorrelation function, while the value of the tangential velocity *v* was obtained from the first data point of the observed mean square displacement 〈*r*^2^〉 of each strain, by assuming ballistic movement at short times, *i.e*. 〈*r*^2^〉 ≈ 〈*v*^2^〉*t*^2^, therefore 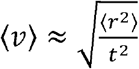.

**S9 Fig: Comparison of motility parameters under different experimental conditions**. (A) Left: Quantification of the number of flagella of individual bacteria grown in TB medium at 30 °C. The flagellation pattern was determined by anti-flagellin immunostaining. The average number of flagella per cell ± s.d. was calculated using Gaussian non-linear regression analysis. Right: Representative fluorescent microscopy images of anti-flagellin immunostaining. The flagellar filament was detected using anti-FliC immunostaining (green) and DNA was stained using DAPI (blue). (B) Measurement of swimming speed *v*, body rotational speed **Ω** and processivity *P* = *v* / **Ω** relative to the wild type for hook-length mutants grown in in LB at 37 °C and TB at 30 °C. (C) Measurement of swimming speed v, body rotational speed **Ω** and processivity *P* = *v* / **Ω** relative to the wild type for chemotactic deficient (Δ*cheY*) hook-length mutants grown in LB at 37 °C and TB at 30 °C.

**S10 Fig: Time dependency of motility parameters for a single dataset**. Absolute values (upper panels) and corresponding data normalized to the wt strain FliK405 (lower panels). (A) Swimming speed *v*, (B) body angular velocity ***Ω***, (C) processivity *P* = *v* / ***Ω***, (D) straightness of trajectory *R*, (E) relative width *S*, of the speed distribution P(v). Mean values presented in Fig 4 are obtained by averaging over the time-window 10 min < *t* < 60 min.

**S11 Fig: Relative width *S* = *σ* / *v* of the speed distribution *P*(*v*) of FliK mutants**.

Upper panel: FliK mutants capable of chemotaxis; Lower panel: smooth swimming FliK mutants due to deletion of *cheY*. Data points represent the mean of an average of up to 5 experiments and were normalized to the wt. Uncontrolled hook-length mutants are shown in red.

**S12 Fig: Representative data set of absolute values obtained from normalized quantities**. (A) Swimming speed *v*, (B) body rotational speed ***Ω***, (C) processivity *P* = *v*/ ***Ω***, (D) straightness of trajectory R, (E) relative width S, of the speed distribution P(v). Upper panel: FliK mutants capable of chemotaxis; Lower panel: smooth swimming FliK mutants due to deletion of *cheY*. Data points represent the mean of an average of up to 5 experiments. Uncontrolled hook-length mutants are shown in red.

**S1 Movie: Motility performance of hook-length variants revealed by single-cell tracking in quasi 2D**. Exemplary movies illustrating the swimming behavior of FliK335, FliK405 (wt), FliK620 and FliK785 are shown.

**S1 Data**: Summary of statistical analysis of the number of flagellar filaments per cell shown in Fig 1D.

**S2 Data**: Numerical data used in Fig 1, Fig 3, Fig 4, S1 Fig, S2 Fig, S3 Fig, S4 Fig, S5 Fig, S7 Fig, S8 Fig, S9 Fig, S10 Fig, S11 Fig, S12 Fig.

**S3 Data**: Numerical data used in Fig 2.

**S4 Data**: Numerical data used in S6 Fig.

